# Biological Aging of CNS-Resident Cells Alters the Clinical Course and Immunopathology of Autoimmune Demyelinating Disease

**DOI:** 10.1101/2021.12.26.473589

**Authors:** Jeffrey R. Atkinson, Andrew D. Jerome, Andrew R. Sas, Ashley Munie, Cankun Wang, Anjun Ma, W. David Arnold, Benjamin M. Segal

**Author notes:** Corresponding Author: Segal, BM.

## Abstract

Biological aging is the strongest factor associated with the clinical phenotype of multiple sclerosis (MS). Relapsing remitting MS (RRMS) typically presents in the third or fourth decade, while the mean age of presentation of progressive MS (pMS) is 45 years old. Here we show that experimental autoimmune encephalomyelitis (EAE), induced by the adoptive transfer of encephalitogenic CD4^+^ Th17 cells, is more severe, and less like to remit, in middle-aged compared with young adult mice. Donor T cells and neutrophils are more abundant, while B cells are relatively sparse, in central nervous system (CNS) infiltrates of the older mice. Experiments with reciprocal bone marrow chimeras demonstrate that radio-resistant, non-hematopoietic cells play a dominant role in shaping age-dependent features of the neuroinflammatory response, as well as the clinical course, during EAE. Reminiscent of pMS, EAE in middle-aged adoptive transfer recipients is characterized by widespread microglial activation. Microglia from older mice express a distinctive transcriptomic profile, suggestive of enhanced chemokine synthesis and antigen presentation. Collectively, our findings suggest that drugs that suppress microglial activation, and acquisition or expression of aging-associated properties, may be beneficial in the treatment of progressive forms of inflammatory demyelinating disease.

## Introduction

Traditionally, people with multiple sclerosis (MS) are categorized into 2 major clinical subsets based on whether they experience a relapsing-remitting or progressive course. Relapsing-remitting MS (RRMS) is characterized by discrete, self-limited episodes of neurological signs and symptoms that persist for weeks to months, followed by a partial or full recovery. Clinical relapses are associated with the appearance of acute inflammatory demyelinating lesions in central nervous system (CNS) white matter, which are driven by the infiltration of lymphocytes and myeloid cells from the blood stream into the CNS across leaky cerebrovascular venules (1). In contrast, progressive MS (pMS) involves an insidious, gradual decline in neurological function (most often in the form of worsening paraparesis, gait imbalance, and/or dementia), in the absence of focal blood-brain-barrier (BBB) breakdown or the influx of peripheral immune cells. Typical pathological features of pMS include slowly expanding white matter lesions, composed of a gliotic core surrounded by a rim of activated microglia, as well as widespread microglia activation in the macroscopically normal appearing white matter (1, 2). Disease modifying therapies (DMT) that are approved for the treatment of MS mostly target lymphocytes in the peripheral immune system. While highly efficacious in RRMS, they have a modest therapeutic impact, at best, in pMS. There is an unmet need for new treatments that slow, or even block, the accumulation of disability in individuals with pMS. The identification of novel therapeutic targets pertinent to pMS will require a deeper understanding of the pathogenic effector cells, mediators, and pathways specific to the manifestation of progressive disease.

Biological aging is the strongest factor associated with MS clinical phenotype. RRMS typically presents in the third or fourth decade, while the mean age of onset of pMS is 45 years old (3). The age of onset of pMS is similar whether it occurs at clinical presentation (referred to as primary progressive MS or PPMS), or follows an initial RR phase (secondary progressive MS or SPMS). Progressive MS is rare before age 40, and virtually non-existent in the pediatric population (4). MS relapse rates tend to decline with disease duration and chronological age (5). Collectively, these data suggest that physiological changes associated with biological aging interact with the autoimmune response that drives CNS damage during MS, in a manner that transforms the clinical course and underlying neuropathology to a progressive pattern. Biological aging could impact the pathogenesis of MS via direct effects on immune cells or CNS-resident cells. Regarding the former, aging leads to immunosenescence, including a chronic, low-grade inflammatory state referred to as “inflamm-aging”. Regarding the latter, aging CNS tissues may be more vulnerable to inflammatory insults and less capable of initiating repair. The relative contributions of these aging-related phenomena to chronic active lesions, widespread microglial activation, and relentless accumulation of disability during pMS remains to be elucidated.

Experimental autoimmune encephalomyelitis (EAE), the most popular animal model of MS, generally presents with an acute monophasic or RR course (6). The pathological features of EAE, such as multifocal BBB breakdown and CNS influx of hematogenous leukocytes, are more reminiscent of RRMS than pMS. EAE is traditionally induced in young adult mice, between 8-12 weeks of age (7). There is a dearth of studies examining the impact of biological aging on animal models of inflammatory demyelination. Here we compare the clinical course, neuropathology, and inflammatory milieu of EAE in young adult versus middle-aged mice.

## Results

### Biological aging exacerbates the effector phase of EAE

In order to distinguish between the effect of biological aging during the T cell-priming and effector stages of EAE, we employed an adoptive transfer model in which IL-23-polarized MOG_35-55_-reactive CD4^+^ T cells from donors of different ages are injected into naïve syngeneic recipients of different ages. Traditionally, EAE is induced in young adult mice, between 8 to 12 weeks of age. For the middle-aged cohort, we used mice 40-44 weeks of age, based on their general developmental similarity to humans at the mean age of onset for pMS (8). We found no significant differences in the clinical course of EAE induced by encephalitogenic CD4^+^ T cells derived from MOG-immunized young adult versus middle-aged donors (Supplementary Figure 1). In contrast, the clinical course of EAE was accelerated and exacerbated in middle-aged, compared with young adult, adoptive transfer recipients that had been injected with the same pool of encephalitogenic CD4^+^ T cells (Fig. 1). Middle-aged recipients developed neurological deficits earlier, had higher peak scores, and more cumulative neurological disability (Fig. 1 A-C). Middle-aged recipients exhibited greater weight loss, and a significant proportion succumbed to disease (58%), while all of the young recipients survived (Fig.1 D, E). Moreover, approximately 80% of young adult recipients underwent clinical remission, compared with approximately 40% of their middle-aged counterparts (Fig. 1 F). This divergence in clinical courses between the 2 groups was reflected in pathological findings. Immunohistochemical analysis of spinal cord sections revealed more extensive demyelination, associated with parenchymal infiltration by CD45^hi^ hematogenous leukocytes, in the middle-aged cohort (Fig. 2). Electrophysiological studies were performed to assess the functional integrity of the spinal cord. Motor-evoked responses following cervical spinal cord stimulation showed delayed responses in all mice at peak and chronic stages of EAE compared with baseline, indicative of demyelination in the corticospinal tracts (Fig. 3B). By week 2 post-transfer, the delay in nerve signal propagation was significantly longer in middle-aged hosts (Fig. 3C), consistent with enhanced myelin damage, as had also been revealed by immunohistochemistry. In addition, from day 13 post-transfer onward, middle-aged mice exhibited a greater reduction in motor-evoked potential amplitudes compared with young adult mice, signifying increased axonal dysfunction and/or loss (Fig. 3D). Motor unit number estimation (MUNE) is calculated by dividing the maximum compound muscle action potential amplitude, obtained with sciatic nerve stimulation, by the average single motor unit potential. MUNE, which reflects the number of motor units innervating single muscles, was lower in the sciatic innervated hindlimb muscles of middle-aged hosts on day 27 post-transfer (Fig. 3E), consistent with a greater loss or dysfunction of motor neurons in the lumbar spinal cord.

**Figure 1.**
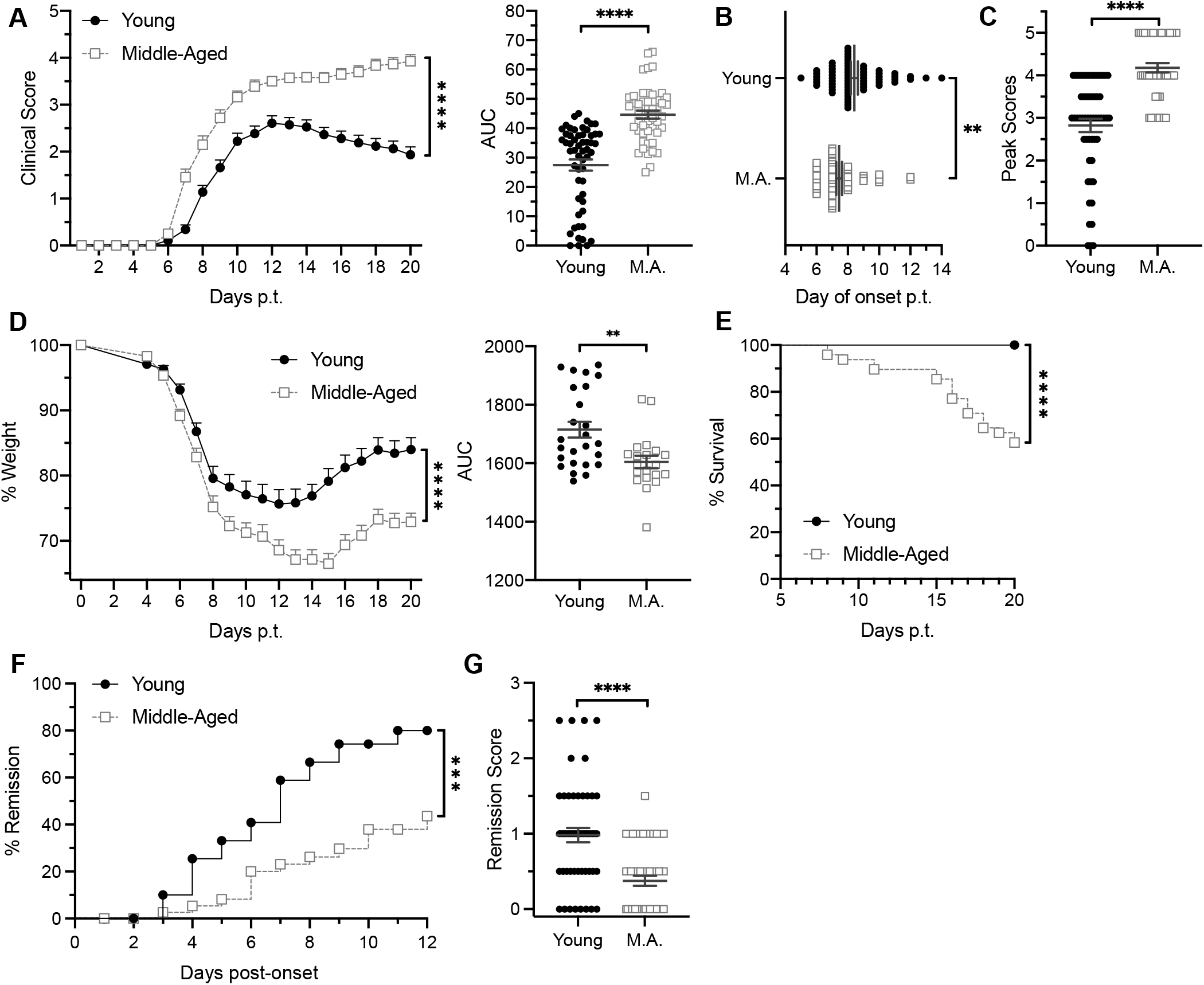
Middle-aged mice exhibit an exacerbated, non-remitting course of Th17-mediated EAE. EAE was induced by the adoptive transfer of myelin-reactive Th17 cells into young or middle-aged mice. (**A-C**) Young (n=70) and middle-aged (n=62) adoptive transfer recipients were scored daily for severity of neurological deficits by an examiner blinded to the identity of experimental groups. Data were pooled from 4 independent experiments. (**A**) Mean clinical scores of young and middle-aged hosts over time (left panel). The area under the curve (AUC) was measured for individual mice in each group (right panel). (**B**) Day of disease onset post-transfer (p.t.) and (**C**) peak clinical disease scores of individual mice. (**D**) Young (n=40) and middle aged (n=36) recipients were weighed on a daily basis. Data were pooled from 3 independent experiments. The % of baseline weight over time, averaged across young and middle-aged recipients (left panel). The area under the curve (AUC) was measured for individual mice in each group (right panel). (**E**) Percent survival over the disease course. (**F**) Percent of mice undergoing remission on each day post clinical onset (left panel). (**H**) Difference between clinical score at peak disease minus score at day 20 post-cell transfer for each mouse. Each symbol in **B** and **C**, and the right panels of **A**, **D**, and **F**, represents data generated from a single mouse. Statistical significance was determined using unpaired 2-tailed Student’s *t* test. Curves in the left panels of **A** and **D** were compared using a mixed effects model. Curves in **F** and **G** were compared using the Log rank test. Error bars indicate mean ± SEM. *p <0.05, **p <0.01, ***p <0.001, ****p <0.0001.

**Figure 2.**
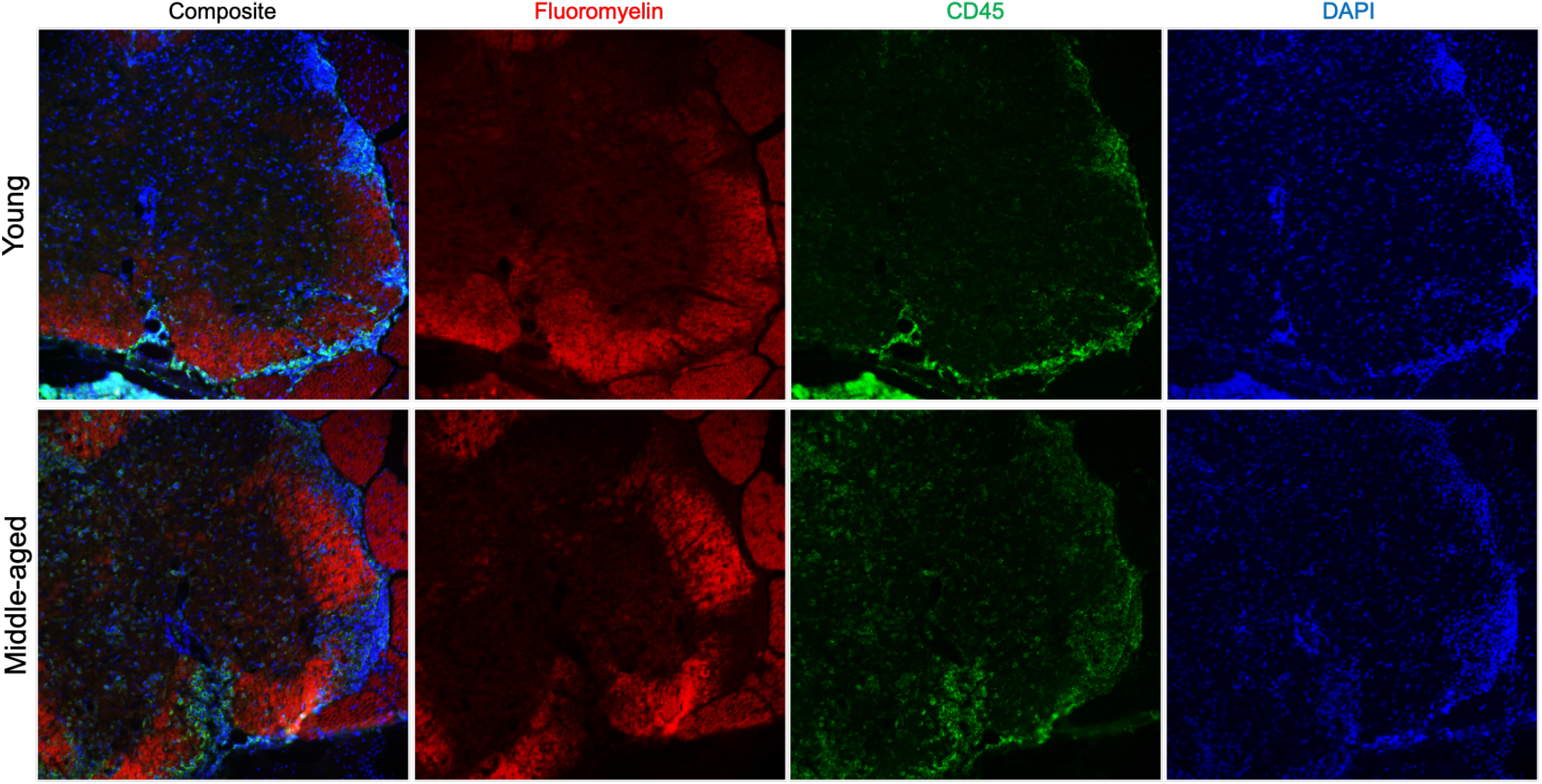
Spinal cord demyelination and inflammatory infiltration are exacerbated in middle-aged recipients. Representative spinal cord sections of young (upper panels) and middle-aged (lower panels) adoptive transfer recipients of encephalitogenic Th17 cells. Sections were stained for myelin (Fluoromyelin; red), CD45 (green), and DAPI (blue). All images were acquired at 20x magnification.

**Figure 3.**
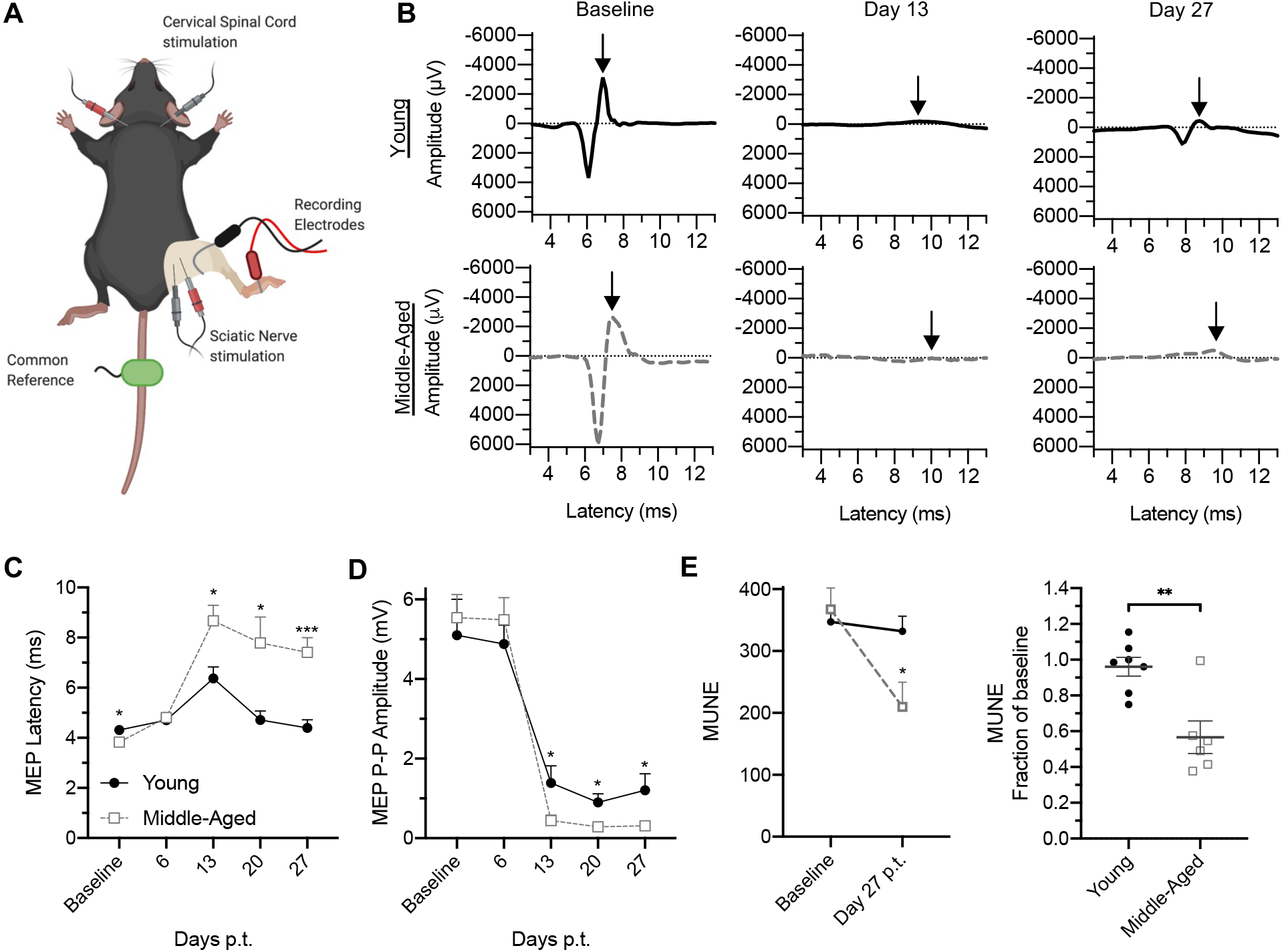
Electrophysiological studies indicate increased demyelination and neuronal/axonal drop out in the spinal cords of middle-aged mice with EAE. (**A-E**) EAE was induced in young adult (n=10) and middle-aged (n=10) mice by the adoptive transfer of encephalitogenic MOG-specific CD4+ T cells. Mice were anaesthetized for electrographic recordings. (**A**) Depiction of electrode placement. (**B**) Representative cervical motor-evoked potentials recorded from the right gastrocnemius following cervical spinal cord stimulation at baseline, day 13, and day 27 post-cell transfer. Arrows indicate negative peaks of the cervical motor-evoked potentials. (**C, D**) Electrographic measurements were obtained at baseline and then on a weekly basis post-transfer (p.t.). Measurements of motor-evoked potential latency (**C**), and peak-to-peak (P-P) amplitude (**D**) were averaged across each group. (**E**) Motor unit number estimation (MUNE) was calculated at baseline and on day 27 post transfer. Mean values for each group are shown (left panel). The ratio of MUNE at day 27 post-cell transfer over baseline is shown for individual mice (right panel). Statistical significance determined by unpaired 2-tailed Student’s *t* test. *p <0.05, **p <0.01, ***p <0.001. Error bars indicate mean ± SEM.

### Encephalitogenic T cells and neutrophils are expanded, while B cells are retracted, in CNS infiltrates of middle-aged mice with EAE

The exacerbated clinical course of EAE in older mice could be secondary to a more virulent neuroimmune response, increased susceptibility of CNS-resident cells/myelin to immune-mediated damage, or a combination of these mechanisms. To investigate the former possibility, we compared the cellular composition of neuroinflammatory infiltrates between middle-aged and young adult mice on day 10 post-T cell transfer. There were significantly higher frequencies of CD4^+^ T cells and neutrophils, and a lower frequency of B cells, among CD45^+^ cells isolated from the spinal cords of the middle-aged recipients (Fig. 4A-C). There were no differences between groups in the frequencies of macrophage/ monocytes or monocyte-derived dendritic cells (Supplementary Figure 2). For our experiments, we routinely obtain MOG_35-55_-primed CD4^+^ T cells from CD45.1^+^ congenic donors, in order to distinguish the transferred encephalitogenic T cells from CD45.2^+^ bystander host T cells. We found that CD45.1^+^ CD4^+^ T cells were disproportionately expanded in the CNS of middle-aged recipients (Fig. 4B). A broad panel of pro-inflammatory and chemotactic factors were up-regulated in spinal cord lysates of all mice with EAE, but were consistently elevated to higher levels in older mice. These factors include the chemokines CCL5, CXCL9 and CXCL10, which have been shown to target encephalitogenic T cells (9, 10), and CXCL1 and CXCL2, the major chemoattractants of mature neutrophils (11, 12) (Fig. 5). Neutrophil-mobilizing factors GM-CSF and G-CSF were also expressed at higher levels in spinal cord lysates obtained from older mice.

**Figure 4.**
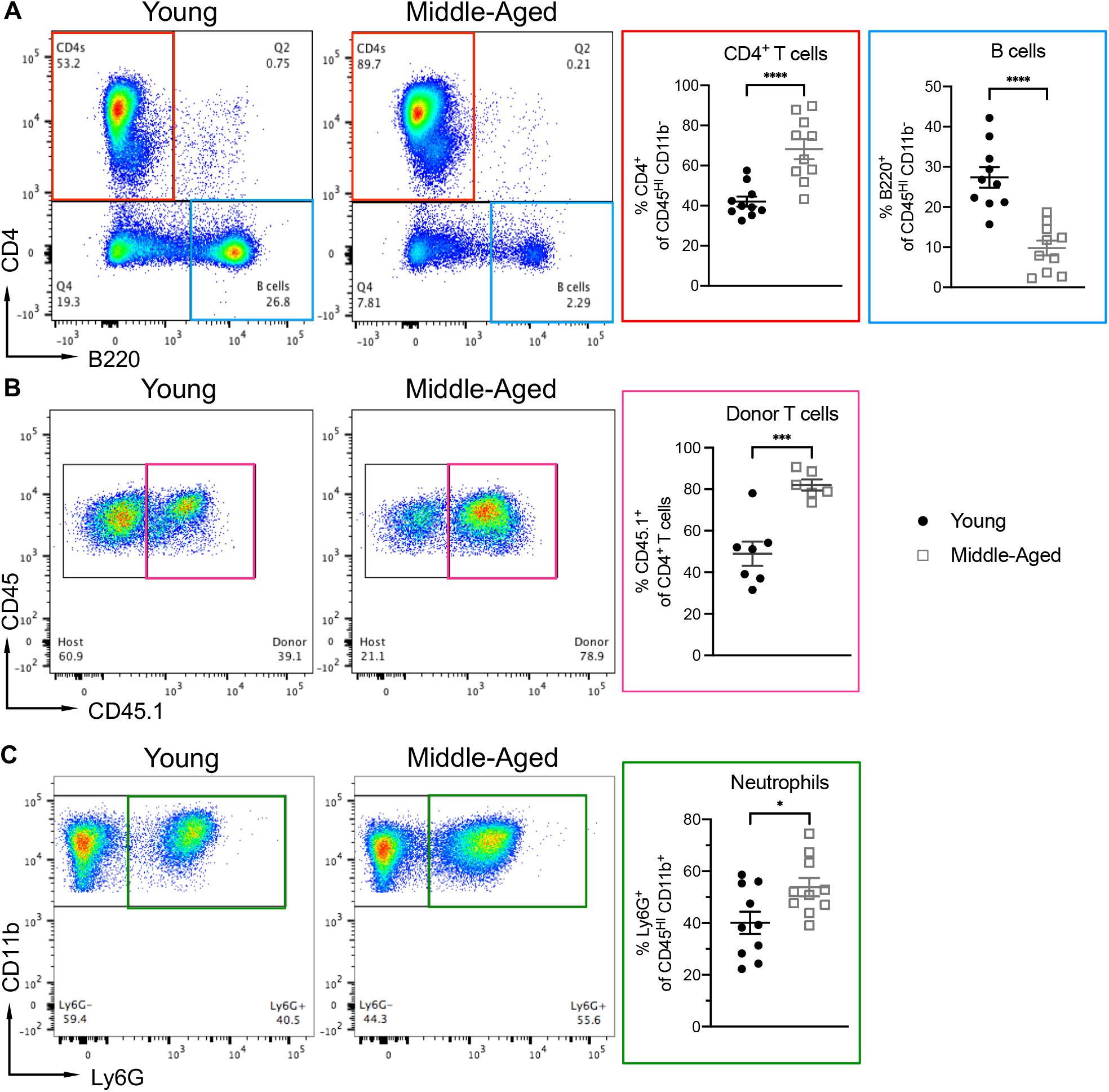
The composition of CNS-infiltrating immune cells differs between middle-aged and young adult mice with EAE. (**A-C**) Mononuclear cells were isolated from the spinal cords of young adult and middle-aged mice on day 10 post-transfer of encephalitogenic CD45.1^+^ CD4^+^ T cells for flow cytometric analysis, gating on all CD45^+^ cells. Representative dot plots (left panels). The frequencies of CD4^+^ T cells and B cells (**A**), CD45.1^+^ donor CD4+ T cells (**B**), and neutrophils (**C**), among CNS-infiltrating CD45^+^ cells (right panels). Each symbol represents an individual mouse. Data were pooled from 3 independent experiments with a total of 7-10 mice per group. Statistical significance was determined using the unpaired 2-tailed Student’s *t* test. *p <0.05, ***p <0.001, ****p <0.0001. Error bars indicate mean ± SEM.

**Figure 5.**
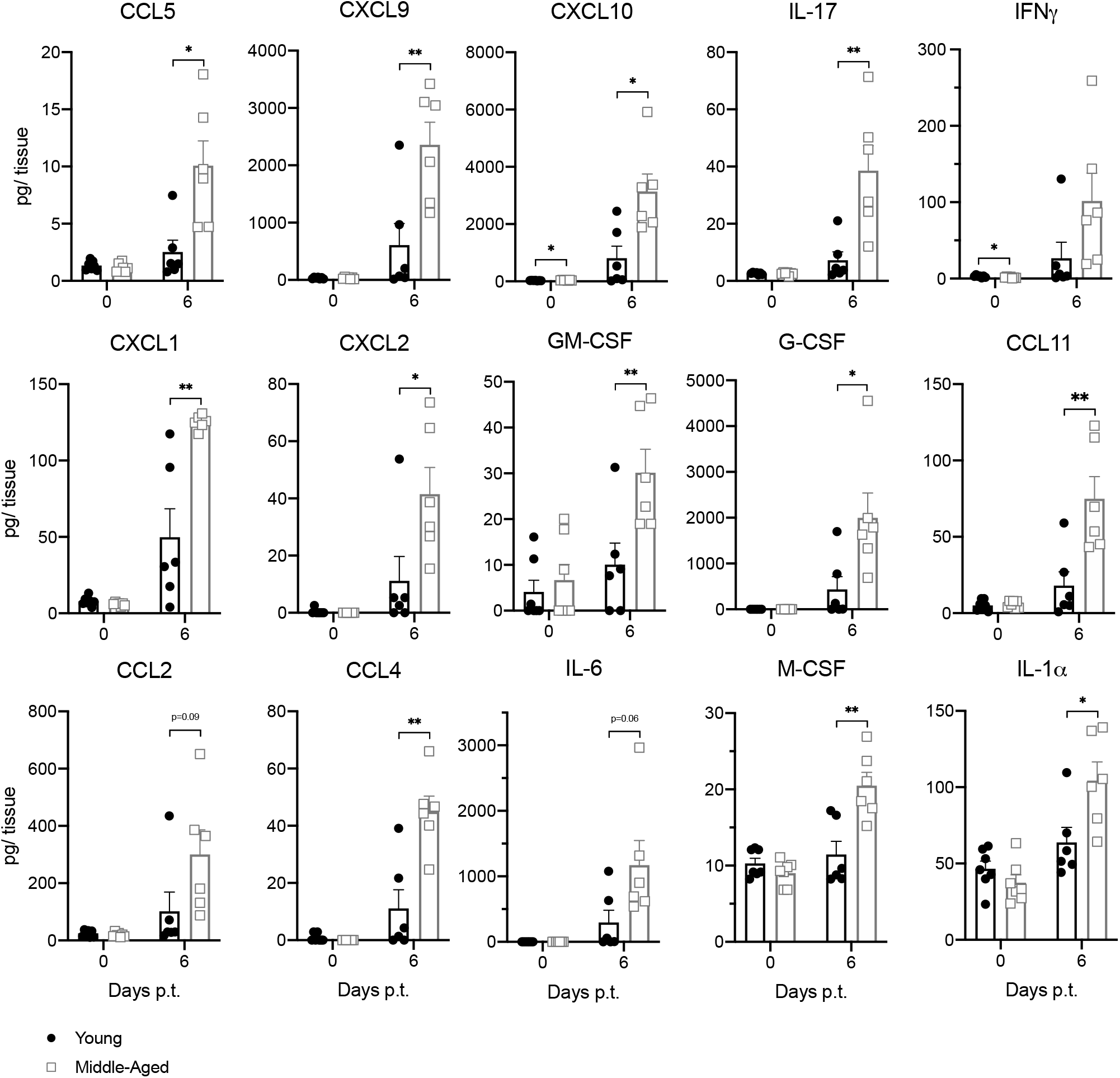
Pro-inflammatory proteins and elevated in the spinal cords of middle-aged mice with adoptive transfer EAE. Spinal cord homogenates were obtained from young (closed circles) and middle-aged (open squares) naïve mice (left bars) or adoptive transfer recipients on day 6 post-transfer (right bars). A panel of pro-inflammatory factors were measured using the Luminex bead-based multiplex platform. Data were pooled from 2-4 independent experiments with a total of 7-13 mice/group. Each symbol represents a single mouse. Statistical significance was determined using the unpaired 2-tailed Student’s *t* test. *p <0.05, ***p <0.001, ****p <0.0001. Error bars indicate mean ± SEM.

### GM-CSF promotes the early, but not late, stage of exacerbated EAE in aged mice

We have previously shown that GM-CSF receptor (GM-CSFR)-deficient mice are relatively resistant to adoptively transferred EAE (13). They experience a milder course than wild-type counterparts, with an increased rate of remission. In addition, GM-CSFR deficiency is associated with a lower percent of CD4^+^ donor T cells and neutrophils, and a higher percent of B cells, in CNS infiltrates, which is the mirror image of the pattern that we observed in middle-age, wild-type mice with EAE. Collectively, these observations suggest that the exacerbated form of EAE that occurs in middle-aged wild-type mice is GM-CSF-driven. Indeed, the absolute number of GM-CSF-expressing donor CD4^+^ T cells in CNS infiltrates, the level of intracellular GM-CSF in CNS donor T cells, and the level of GM-CSF protein in CNS lysates, were higher in the older versus younger mice with EAE (Fig. 5, Fig.6A, B). Furthermore, single-cell RNA sequencing revealed heightened expression of transcripts encoding GM-CSF, as well as IL-17 and other pro-inflammatory factors, in CD4^+^ T cells isolated from the CNS of middle-aged mice on day 6 post-transfer, when compared with their younger counterparts (Fig. 6C). Treatment of middle-aged hosts with a neutralizing antibody against GM-CSF, beginning from the day of transfer onward, delayed clinical onset and ameliorated the early clinical course (Fig. 6D, E). However, mean peak scores, cumulative chronic disability, and mortality rates were similar between the anti-GM-CSF and control antibody treatment groups (Fig. 6D, F, and data not shown). Postponing the initiation of anti-GM-CSF treatment to the time of peak disease had no therapeutic impact (data not shown). Based on these data, we concluded that GM-CSF promotes aggressive EAE in older recipients in the early phase of disease, but becomes dispensable as the disease progresses towards a more chronic phase.

**Figure 6.**
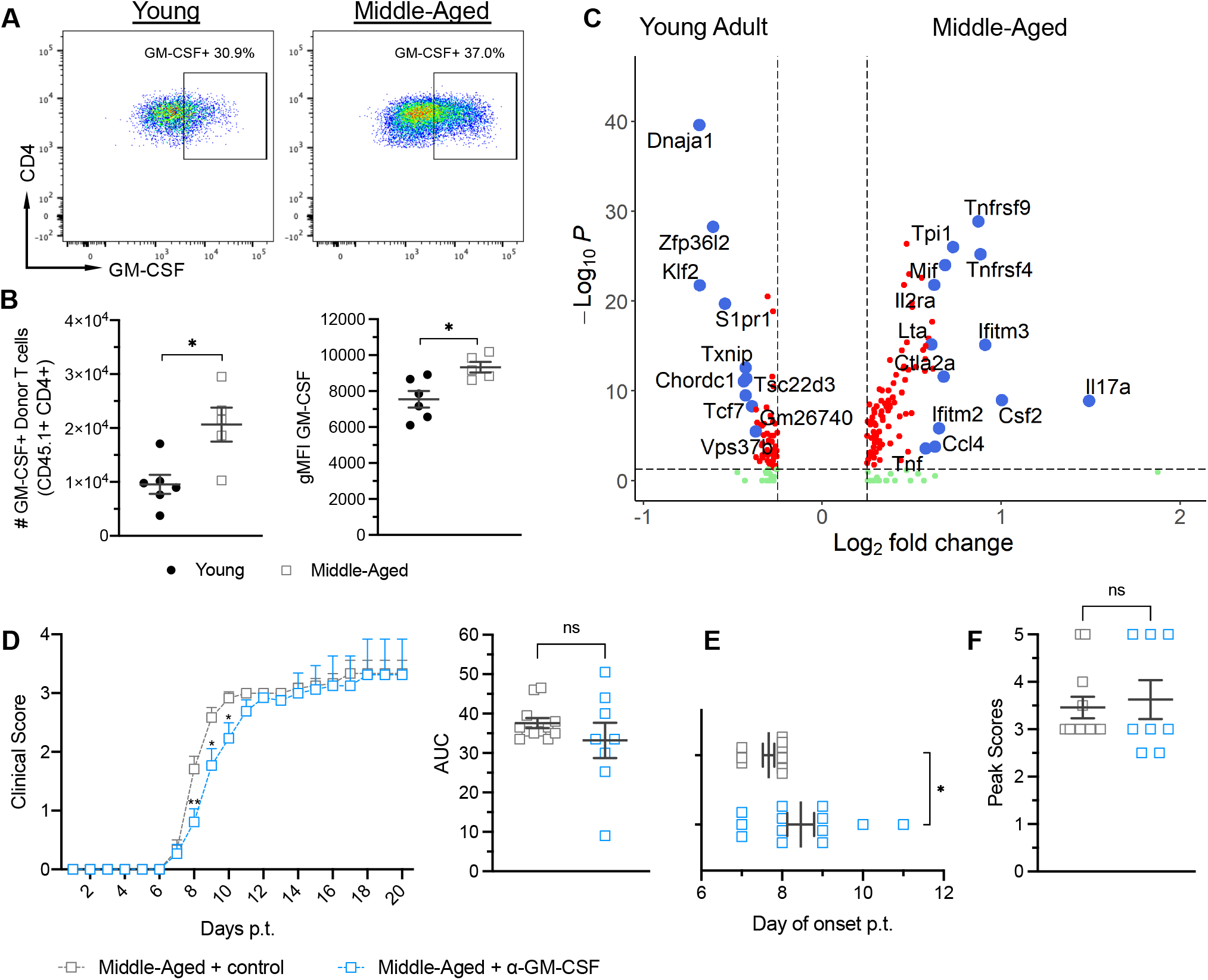
GM-CSF promotes exacerbated EAE in middle-aged mice early, but not late, in the clinical course. (**A, B**) CNS mononuclear cells were harvested from the spinal cords of young adult and middle-aged mice on day 10 post-T cell transfer for analysis by flow cytometry. (**A**) Representative dot plots showing intracellular GM-CSF expression, gating on CD45.1^+^ donor CD4^+^ T cells. (**B**) Total number of GM-CSF^+^ donor T cells (left panel), and geometric mean fluorescent intensity (gMFI) of GM-CSF in donor T cells (right panel), isolated from the spinal cords of individual mice. Data were pooled from 2 independent experiments with 5-6 mice/group. (**C**) CD45^+^ mononuclear cells, FACS sorted from young adult or middle-aged mice with EAE on day 6 post-cell transfer, were analyzed by scRNAseq. Gene expression changes in CNS CD4+ T cells, comparing young and middle aged mice, are shown in a volcano plot. (**D-F**) Middle-aged mice were injected i.p. with anti-GM-CSF neutralizing antibody or control antibody every other day from the time of encephalitogenic T cell transfer onward (n= 14 mice/group; pooled from 2 independent experiments). Mice were scored for severity of neurological deficits on a daily basis by raters blinded to the identity of the experimental groups. (**D**) Mean clinical scores of mice in each group over time (left panel). Area under the curves (AUC) of individual mice (right panel). (**E**) Day of clinical disease onset post-transfer and (**F**) peak clinical disease scores of individual mice. Statistical significance was determined using the unpaired 2-tailed Student’s *t* test. *p <0.05 or as indicated. Error bars indicate mean ± SEM.

### Radio-resistant, non-hematopoietic cells shape the neuroinflammatory infiltrate and drive exacerbated EAE in middle-aged mice

Next, we sought to determine whether the increased susceptibility of older mice to EAE is secondary to the biological aging of radio-sensitive, hematopoietic cells (that are recruited from the circulation to the CNS) and/or radio-resistant, non-hematopoietic cells (including CNS-resident cells). To that end, we constructed reciprocal bone marrow (BM) chimeric mice with young adult or middle-aged BM cell donors and/or irradiated hosts (Fig. 7A). Middle-aged → middle-aged and young adult → young adult chimeras served as controls. EAE was induced in all immune reconstituted chimeric mice via the adoptive transfer of the same pool of MOG_35-55_ encephalitogenic CD4^+^ T cells. Irrespective of the age of the BM cell donors, middle -aged BM cell hosts experienced a severe clinical course of EAE, with high mortality rates, while young adult BM cell hosts experienced a milder course with relatively low mortality rates (Fig. 7 B, C). Furthermore, the cellular composition of neuroinflammatory infiltrates in chimeras correlated with the age of the BM cell host, but not the donor, mouse. Middle-aged BM cell hosts reconstituted with young BM cells contained higher frequencies of CD4^+^ T cells in EAE lesions compared with chimeras that were constructed using young adult hosts, mimicking the results we obtained with non-chimeric mice (Fig. 7D, left panel). Furthermore, the frequency of B cells tended to be lower, while that of neutrophils tended to be higher, in CNS infiltrates of the middle-aged hosts, though not reaching statistical significance (Fig. 7D, middle and right panels). Hence, the age of radio-resistant, non-hematopoietic cells is a dominant factor in determining the phenotype and severity of EAE.

**Figure 7.**
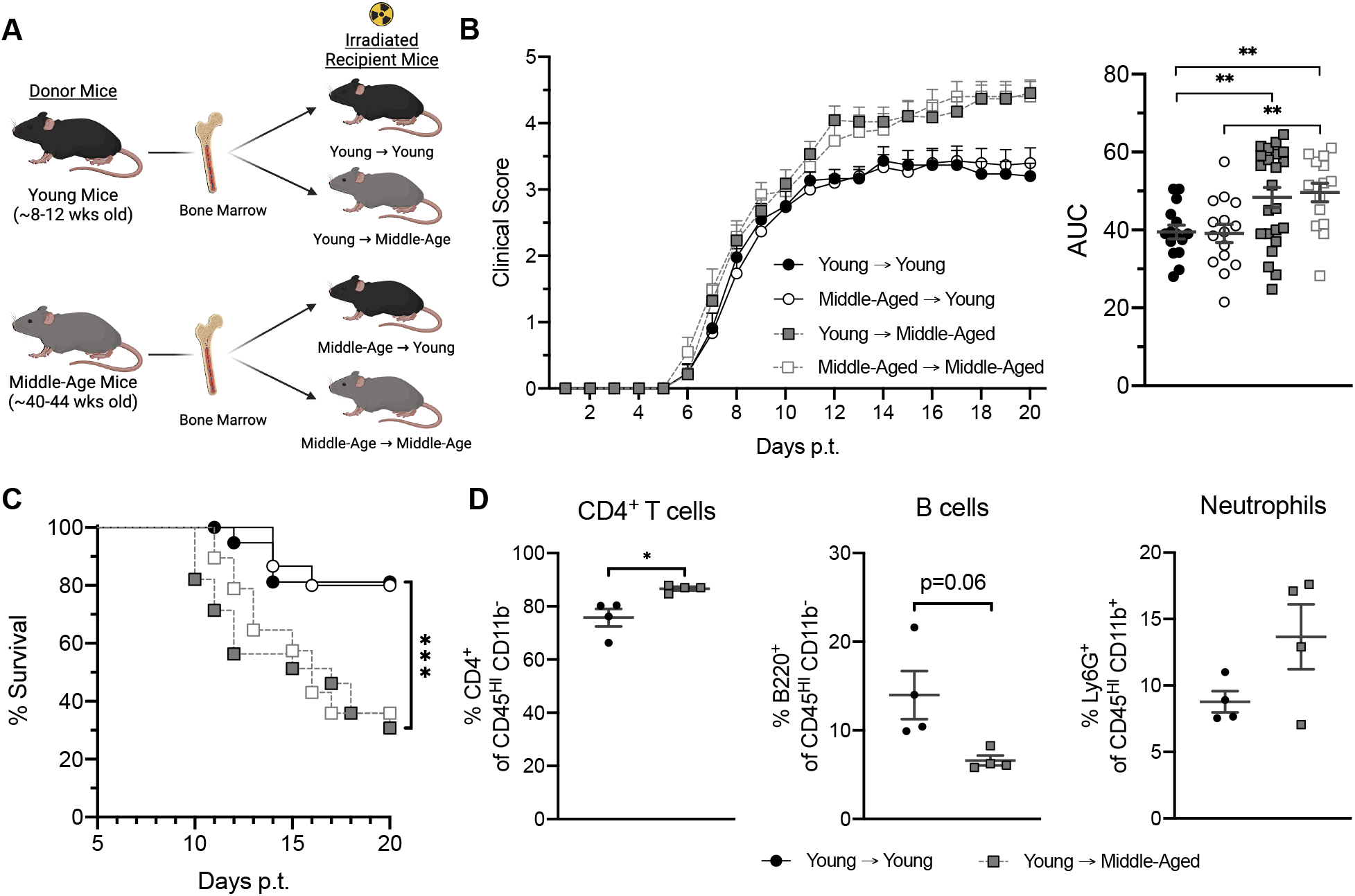
Radio-resistant, non-hematopoietic cells drive exacerbated EAE in middle-aged mice. (**A-D**) Reciprocal bone marrow chimeric mice were constructed with young adult or middle-aged donors and hosts. Following reconstitution, all chimeric mice were injected i.p. with 3 x 10^6^ MOG-primed CD4^+^ Th17 cells. Mice were scored for severity of neurological deficits on a daily basis by raters blinded to the identity of the experimental groups. (**A**) Schematic depicting the construction of reciprocal bone marrow chimeras. (**B**) Mean clinical scores over time (left panel) and area under the curves (AUC) of individual mice (right panel). (**C**) Percent of surviving mice in each group over time. (**D**) Frequencies of CD4^+^ T cells, B cells, and neutrophils among CD45^+^ spinal cord mononuclear cells harvested from individual mice on day 10 post-transfer. Data in (**B**) and (**C**) were pooled from 3 independent experiments with a total of 15-28 mice per group. Each symbol in B, right panel, and in D represent an individual mouse, Data in (**D**) are representative experiment of 3 independent experiments (n=4 mice/group). Statistical significance was determined using the unpaired 2-tailed Student’s *t* test. Curves in **C** were compared using the Log rank test. *p <0.05, **p <0.01, ***p <0.001 or as indicated. Error bars indicate mean ± SEM.

### Aged microglia exhibit distinct transcriptomes and phenotypes during homeostasis, as well as EAE

Our finding that radio-resistant cells influence the make-up of neuroinflammatory infiltrates directed our attention towards the role of microglia. In addition to their potential to serve as antigen-presenting cells, microglia produce chemotactic and pro-inflammatory molecules that could orchestrate the recruitment, positioning, and polarization of leukocyte subsets in EAE lesions. A growing body of data indicates that microglia become spontaneously activated and acquire pro-inflammatory signatures with age (14, 15). In support of these published studies, we found that a significant percent of microglia in naïve middle-aged, but not young adult, mice exhibit enhanced expression of the cell surface marker CD11c (Fig. 8A). Bulk RNA sequencing studies showed that microglia isolated from unmanipulated middle-aged mice express elevated levels of pro-inflammatory genes (Fig. 8B, left upper panel), and lower levels of homeostatic genes (Fig. 8B, left lower panel), compared with microglia from their younger counterparts. Furthermore, a number of genes that were disproportionately up-regulated in the middle-aged microglia (such as *Spp1, Cxcl10, Ccl3, Il1β, Ccl4,* and *Tnf*) overlapped with previously published transcriptomes of aging microglial subsets enriched in geriatric mice (Fig. 8B, right) (14).

**Figure 8.**
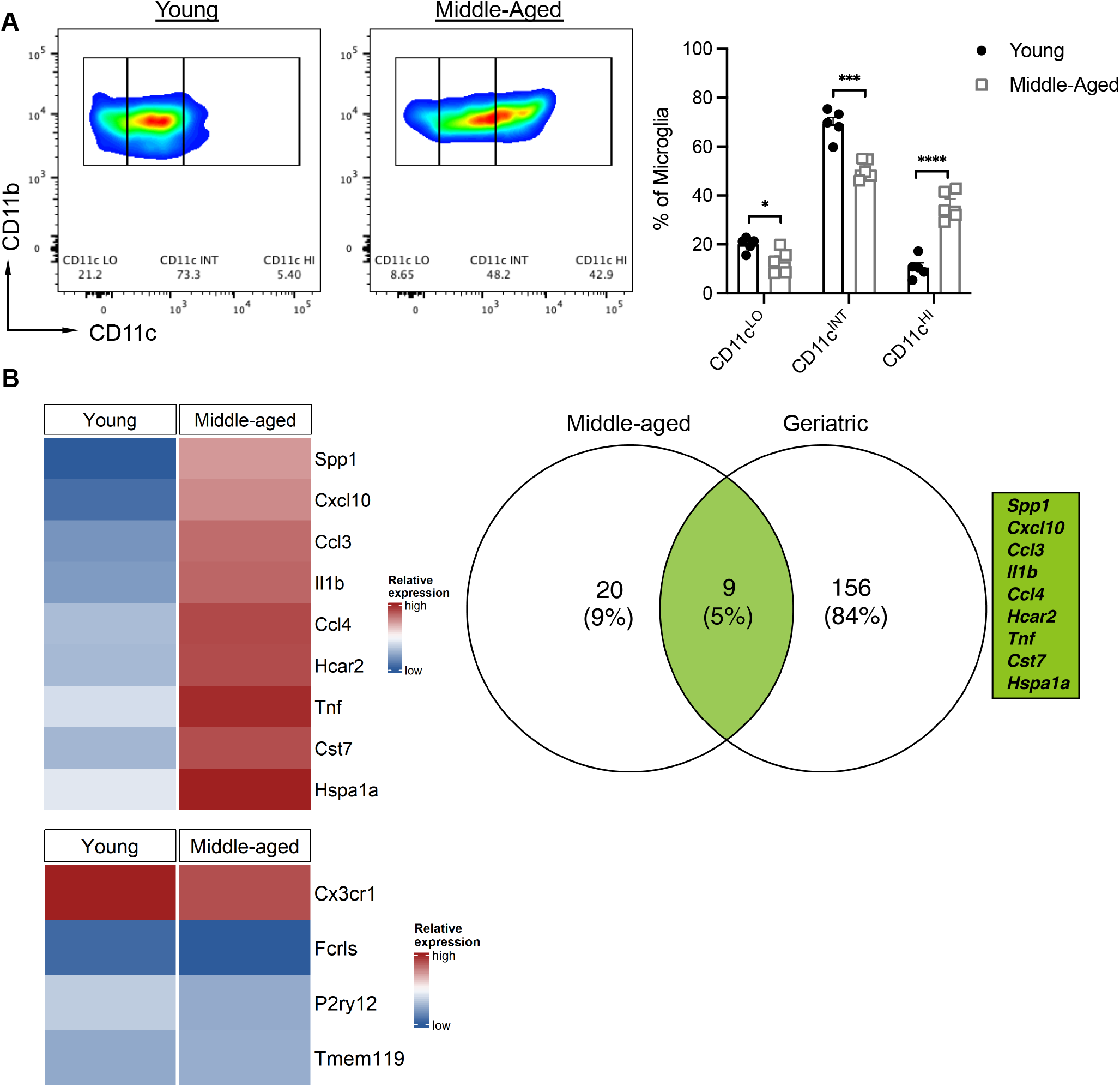
Microglia in middle-aged mice up-regulate genes indicative of a reactive, age-associated phenotype. Mononuclear cells were harvested from the spinal cords of naïve young adult and middle-aged mice. (**A**) Representative dot plots (left and middle panel), and frequencies of CD11c^lo^, CD11c^int^ and CD11c^hi^ cells among CD45^int^CD11b+ microglia (right panel), are shown. Each symbol in the right panel represents an individual mouse. Data were pooled from 2 independent experiments with n=5 mice/group. Statistical significance was determined using the unpaired 2-tailed Student’s *t* test. *p <0.05, ***p <0.001, ****p <0.0001. Error bars indicate mean ± SEM. (**B**) CD45^INT^ CD11b+ cells were FACS sorted and RNA was extracted for bulk sequencing. Venn diagram depicts the number of overlapping genes and overall percent of genes up-regulated in naïve microglia isolated from middle-aged mice compared to geriatric mice (age P540) (left). Heat maps illustrating the relative expression of overlapping genes (right) and homeostatic genes (bottom).

Next, we performed single-cell RNA sequencing (scRNA-seq) of CNS CD45^+^ mononuclear cells isolated from young adult and middle-aged mice during peak EAE. The raw scRNA-seq expression matrix contains 38,057 cells (including 19,899 cells from middle-aged mice and 18,158 cells from young adult) and 19,449 genes. The microglial cells fell into 2 clusters (Fig. 9A). Cluster 1 microglia from middle-aged mice are enriched in transcripts associated with antigen-processing and presentation (including *H2-Aa, H2-Ab1, H2-D1, H-2-Eb1, H2K1, H-2Q7, B2m, Cd74*), and proteasome assembly and function (*Psmb4, Psmb8, Psmb9, Psmb10, Psme1, Psme2*), as well as transcripts that encode pro-inflammatory molecules (*Tnf*, *Il1a, Il1b, Il18)* (Fig. 9 B, C and Fig. 10A), when compared with their counterparts from young adoptive transfer recipients. In contrast, they are relatively deficient in transcripts associated with homeostasis (*Tmem119, Fcrls, Cx3cr1, P2ry12)* and heat shock protein responses (*Dnaja1, Dnajb1, Hspa1a, Hspa1b, Hspa5, Hsp90aa1, Hsp90ab1)* (Fig. 9 B, C and Fig.10A). Cluster 2 microglia from middle-aged mice express relatively high levels of genes encoding chemotactic factors (including *Ccl4, Cxcl2, Cxcl9,* and *Cxcl10)* and relatively low levels of genes associated with homeostasis (including *Fcrls, Tmem119, P2ry12,* and *Cx3cr1)* (Fig. 10 B). Middle-aged microglia in both clusters expressed relatively high expression levels of interferon responses genes (including *Gbp2, Ifitm3, Ifi27l2a, Isg15)* (Fig. 10 A, B), as well as genes that are expressed at high levels in microglia located at the rims of slow expanding lesions in people with pMS (*C1qa, C1qb, C1qc, Cstb, Fth1, Ftl1*, and a panel of ribosomal proteins) (Supplementary Fig. 3).

**Figure. 9.**
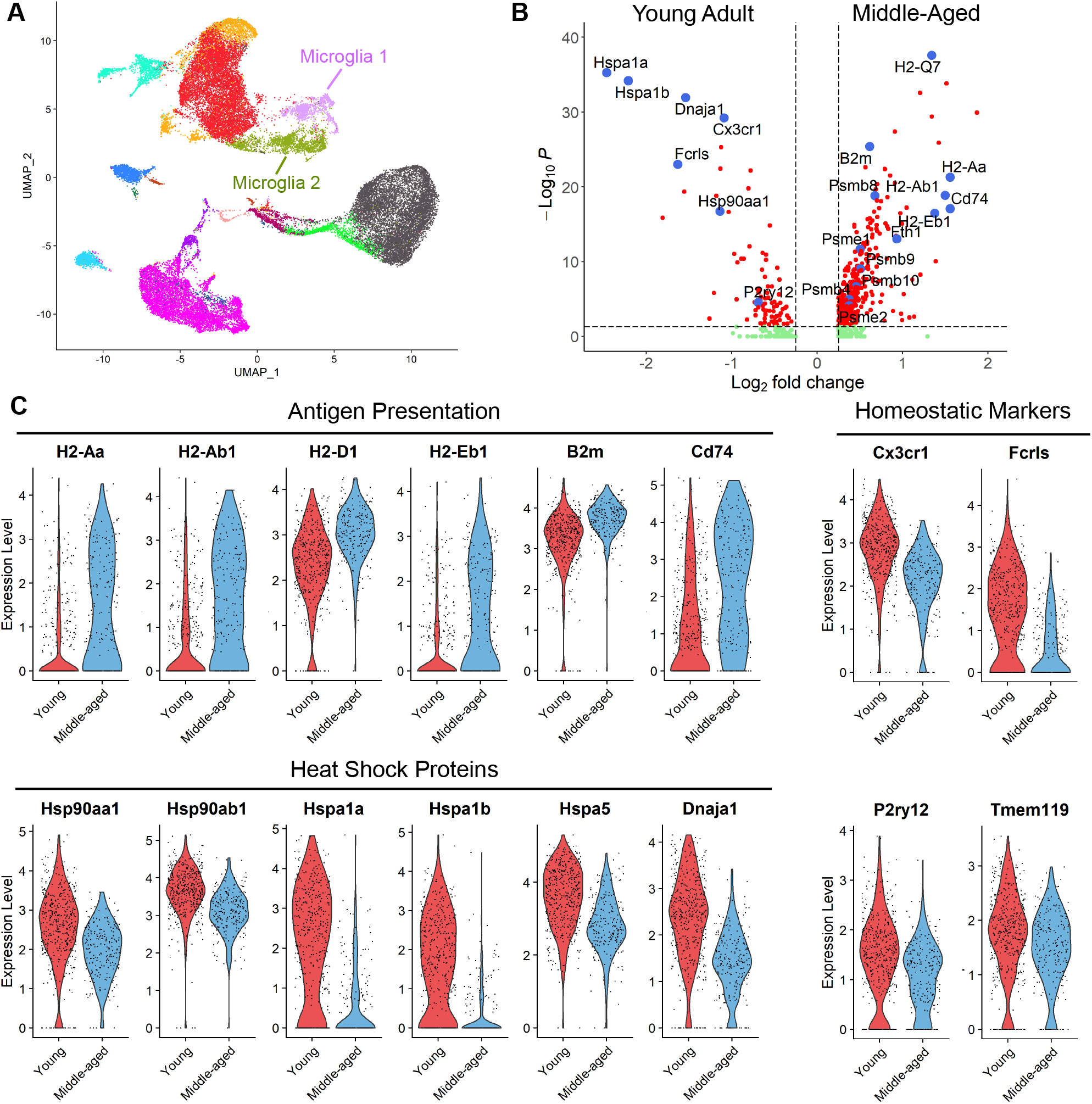
Microglia from middle-aged mice with EAE express distinct transcriptomic signatures. CD45^+^ mononuclear cells were FACS sorted from young adult or middle aged mice with EAE on day 6 post-cell transfer and subjected to scRNA-seq. (**A**) Unified manifold approximation and projection (UMAP) plot showing clustering by cell type, labeled on the basis of known lineage markers. **(B**) Gene expression changes in cluster 1 microglia isolated from young adult versus middle-aged mice, shown as a volcano plot. (**C**) Selected differentially expressed genes in cluster 1 microglia from young adult (red) and middle-aged (blue) adoptive transfer recipients, shown as violin pots. Numbers on the y-axis correspond to the z score for each gene.

**Figure 10.**
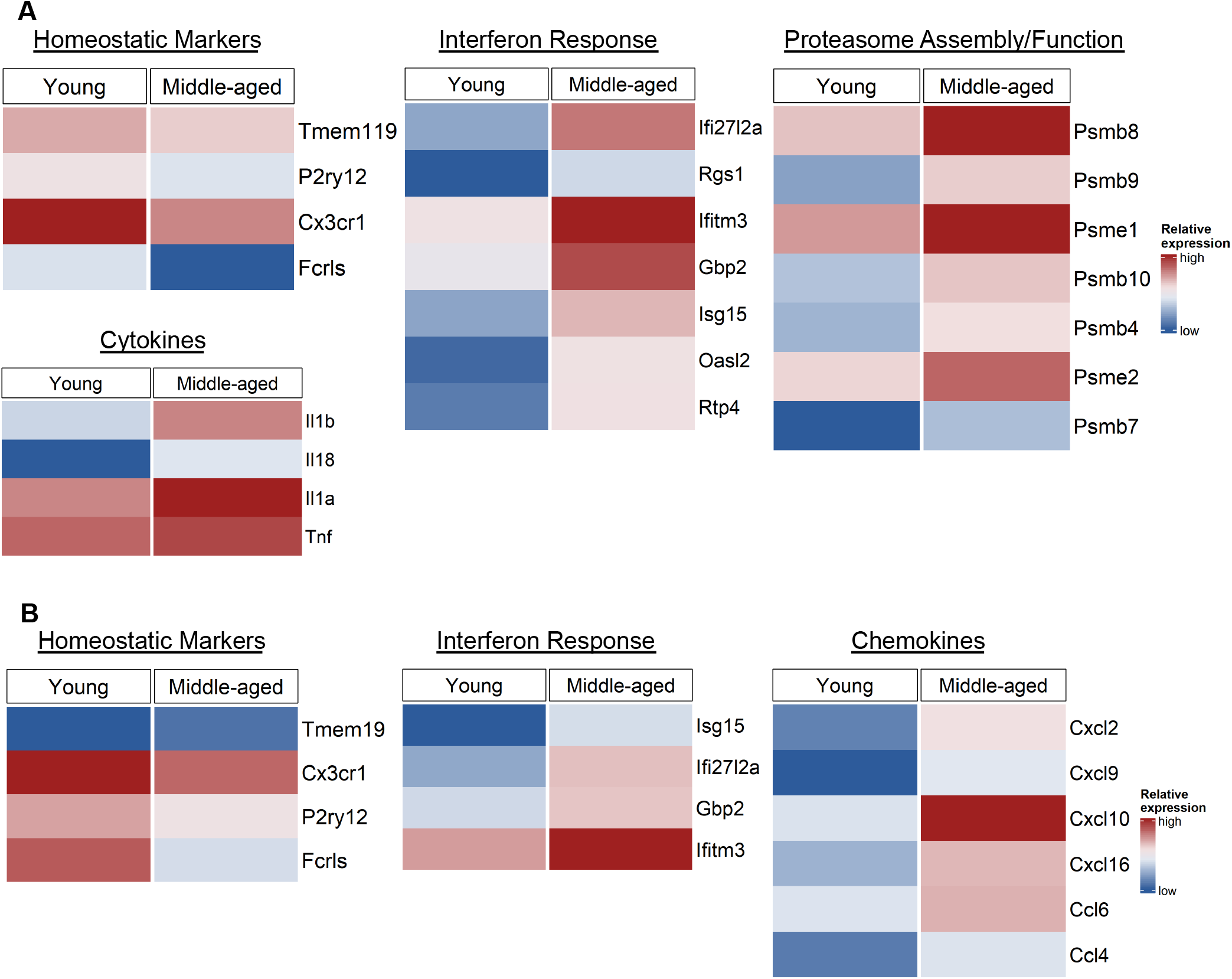
Microglia from middle-aged mice with EAE down-regulate homeostatic genes and up-regulate pro-inflammatory and IFN response genes. Single-cell RNA-sequencing was performed on CD45^+^ CNS mononuclear cells isolated from young adult and middle-aged adoptive transfer recipients as described in the legend to Figure 9. The heat maps show z-score transformed normalized expression values of selected genes in cluster 1 microglia (**A**) or cluster 2 microglia (**B**), comparing young adult and middle-aged recipient mice.

## Discussion

Progressive MS typically presents during middle age, whether or not it is preceded by a RR course. It rarely, if ever, occurs in the pediatric MS population (4). Progressive MS has distinctive pathological features that are not characteristic of RRMS in younger individuals, including widespread microglial activation (1). These observations indicate that biological aging has a profound influence on the evolution and manifestation of autoimmune neuroinflammation, but the underlying mechanisms are poorly understood. Here we show that EAE, induced by the transfer of a common pool of IL-23-polarized, MOG-reactive CD4^+^ T cells, is more severe, and less likely to remit, in middle-aged compared with young adult recipients. We employed an adoptive transfer model in order to distinguish between age-dependent environmental factors in the recipient mouse, as opposed to intrinsic properties of encephalitogenic T cells, that might contribute to the differences in EAE phenotype between the cohorts. The heightened, refractory disability exhibited by older adoptive transfer recipients may reflect an increased vulnerability of aged oligodendrocytes, myelin and/or axons to immune-mediated damage. However, our data indicate that aging has an impact on neuroinflammation itself, since the cellular composition of CD45^+^ cells in the CNS at peak EAE differs markedly between young adult and middle-aged recipients. Donor T cells and neutrophils were consistently more abundant, while B cells were relatively sparse, in the CNS of older mice. Our experiments with reciprocal bone marrow chimeric mice indicate that radio-resistant, non-hematopoietic cells play a dominant role in shaping age-dependent features of the neuroinflammatory response, as well as the clinical course, during EAE. Among radio-resistant host cells, glia are strong candidates for regulators of CNS autoimmunity.

We found that murine microglia spontaneously up-regulate CD11c as they age. This is consistent with previous reports that aging microglia exhibit morphological, phenotypic, and transcriptomic changes indicative of a predisposition towards an activated or pro-inflammatory state (16–18). A number of the pro-inflammatory genes that we identified as upregulated in naive middle-aged, compared with young adult, microglia have previously been reported to be upregulated in microglia from geriatric mice (14). Our working hypothesis is that the increased susceptibility of middle-aged mice to encephalitogenic T cell accumulation, white matter damage, and exacerbated clinical EAE, is driven, at least in part, by aging microglia. Indeed, enhanced microglial reactivity in the aging monkey brain correlates with an increase in perivascular T cell infiltrates in the white matter parenchyma, as well as cognitive impairment (19). Interestingly, genes we found to be highly expressed by microglia from middle aged adoptive transfer recipients are also expressed at high levels by microglia located in the rims of chronic active lesions during pMS, including genes involved in expression of the MHC-II protein complex, ferritin complex, and complement cascade (Supplementary Fig. 3).

The mechanisms underlying age-related changes in microglia are likely multifold. Neuronal maintenance of microglial homeostasis via CX3CL1/CX3CR1 and CD200/CD200R interactions wanes with aging (20–22). In addition, microglia can be stimulated by CNS-penetrant, pro-inflammatory factors, such as IL-6, CCL11, and IL-1β, that are systemically released by peripheral immune cells in the context of “inflamm-aging”, and/or by microbiome-derived molecules, such as LPS, that rise in the bloodstream consequent to age-related increases in gastrointestinal permeability (17, 18, 23–25). Microbial metabolites, which are altered during dysbiosis, also modulate microglial function in an age-dependent fashion. In young adults, gut microbiome-derived tryptophan metabolites and short chain fatty acids (SCFA) cross the blood-brain-barrier and constitutively suppress microglia, thereby reducing the risk of neuroinflammation (26, 27). Dietary supplementation with the SCFA propionic acid is associated with less inflammatory activity, disability progression, and brain atrophy in MS (28). However, production of SCFAs and tryptophan metabolites by gut bacteria declines with aging (29), thereby removing another check on microglial activation.

Single-cell RNA sequencing of CD45^+^ leukocytes, harvested from the CNS at peak disease, revealed 2 clusters of EAE-associated microglia, both of which exhibit dynamic, age-dependent transcriptomic signatures (Fig. 9). By comparison to their counterparts harvested from young adult adoptive transfer recipients, cluster 1 microglia from middle-aged recipients are highly enriched in transcripts that encode molecules engaged in the presentation of antigen to T cells. This finding led us to hypothesize that more efficient antigen presentation by cluster 1 microglia might boost the proliferation of donor T cells, resulting in higher T cell frequencies within the CNS infiltrates of middle-aged mice. However, contrary to that hypothesis, we found comparable frequencies of Ki67^+^ donor T cells among CD45^+^ CNS mononuclear cells isolated from young adult or middle-aged mice at peak EAE (data not shown). Furthermore, CD45^int^CD11b^+^ microglia, purified from young adult or middle-aged mice at peak EAE, stimulated the proliferation of 2D2 cells (T cell receptor transgenic CD4^+^ cells specific for MOG peptide) to a similar degree (data not shown). To further investigate this issue, in future experiments we will compare the proliferation of donor T cells, and the antigen-presenting/T cell-polarizing capacity of sharply-defined microglial subsets, in young adult versus middle-aged hosts at multiple time-points. Interestingly, some of the transcripts upregulated in cluster 1 microglia from middle-aged mice are involved in the assembly and function of proteasomes and processing of MHC Class I restricted peptides. Although our model of EAE is CD4^+^ T cell-driven, CD8^+^ T cells are more prevalent than CD4^+^ T cells in human MS lesions, including in the chronic active lesions that are typical of pMS (30–32). The emergence of microglia particularly well-equipped to activate CD8^+^ T cells might be relevant to the pathogenesis of pMS.

In middle-aged hosts, cluster 2 microglia are enriched in transcripts that encode a range of chemokines including CXCL2 and CXCL10. We and others previously showed that neutrophil migration to the CNS during EAE is dependent on CXCR2-binding chemokines (such as CXCL1, CXCL2)(11, 12, 33, 34), while CD4^+^ T cell migration is dependent on CXCR3-binding chemokines (such as CXCL9, CXCL10 and CXCL11) (9). Elevated production of CXCL2 and CXCL10 by aged cluster 2 microglia might trigger preferential accumulation of donor T cells and neutrophils locally, and thereby underlie the relative enrichment of those subpopulations in the CNS of middle-aged hosts during EAE. The translational significance of this finding is underscored by the observations that pMS patients have a relatively high percentage of circulating CXCR3^+^ lymphocytes (35), and CXCR3^+^ lymphocytes preferentially accumulate in MS lesions (36). Furthermore, CXCL9 and CXCL10 are up-regulated in pMS brains^33^, and expression of CXCR3 on circulating CD8^+^ T cells correlates with MS lesion volume (37). There is also evidence that neutrophil-related factors and chemokines are dysregulated in pMS and correlate with clinical, as well as radiological, measures of CNS tissue damage (25).

In addition to sculpting the neuroinflammatory response via antigen presentation and chemokine production, aging microglia could modulate EAE by releasing soluble factors that directly damage oligodendrocytes and/or axons, such as reactive oxygen and nitrogen species, enzymes and TNF family members (38, 39), and/or that drive the polarization of neurotoxic astrocytes, such as IL-1α and C1q (27, 40). Conversely, we found that aging microglia in cluster 1 downregulate expression of transcripts encoding heat shock proteins 90 and 70, which have been implicated in neuroprotection and the suppression of neurotoxic astrocytes, respectively (41, 42). Increased susceptibility of oligodendrocytes and neurons/axons to immune-mediated insults, combined with impairment of microglial phagocytosis, might escalate CNS deposition of danger-associated molecular patterns (DAMPs) and cellular debris in middle-aged hosts. DAMPs and cellular debris incite the activation of microglia and infiltrating leukocytes, which could fuel a self-amplifying cycle of tissue destruction. Although the adoptive transfer model of EAE in middle-aged mice lacks a number of salient pathological features of pMS (i.e. slowly expanding, chronic active lesions and cortical lesions), it may be useful for investigating how biological aging alters inflammatory demyelinating disease in other ways that simulate progressive MS (i.e. diffuse microglial activation and a non-remitting clinical course), and for testing the efficacy of senolytic drugs, microglial suppressors, or other novel therapeutic interventions potentially beneficial in the pMS subpopulation.

## Methods

### Mice

CD45.1 congenic and wild-type C57BL/6 mice were obtained from Charles River Laboratories or the National Cancer Institute. Mice were housed in micro-isolator cages under specific pathogen-free conditions at the Ohio State University. Both male and female mice were used in experiments.

### Induction and assessment of EAE

8-12 week old mice were immunized subcutaneously with an emulsion consisting of 100 μg of MOG_35-55_ peptide (MEVGWYRSP-FSRVVHLYRNGK; Biosynthesis) emulsified in Complete Freund’s Adjuvant (CFA; Difco), at four sites over the flanks. Inguinal, axial, and brachial lymph nodes and spleens were harvested from donor mice 10-14 days post-immunization and passed through a 70 μm strainer (Fisher Scientific) to obtain a single-cell suspension. The cells were cultured for 96 hours with MOG_35-55_ peptide (50 μg/mL) and Th17-polarizing factors as follows: recombinant murine (rm)IL-23 (8 ng/ml; R&D Systems), rmIL-1α (10 ng/ml; Peprotech) and anti–IFNγ (10 μg/ml; clone XMG1.2, Bio X Cell). After 96 hours, the cells were harvested, washed and resuspended in fresh media. CD4^+^ T cells were purified via positive-selection, magnetic-activated cell sorting (MACS) using L3T4 magnetic microbeads (Miltenyi Biotec) per the manufacturer’s protocol. CD4^+^ T cells (90-98% purity) were transferred via intraperitoneal (i.p.) injection to naïve C57BL/6 recipients. Recipient mice were weighed and observed daily for clinical disability and rated using a 5-point scale as described previously. Briefly, 0.5, partial tail paralysis; 1, full tail paralysis; 1.5, hindlimb weakness demonstrated by ability to correct from a prone position in <1s; 2, hindlimb weakness demonstrated by ability to correct from a prone position in >1s; 2.5, hindlimb weakness demonstrated by severe gait abnormality; 3, partial hindlimb paralysis demonstrated by the inability to elevate hindquarters; 3.5 complete paralysis in one hindlimb; 4, complete hindlimb paralysis; 4.5, moribund; and 5, dead.

### CNS mononuclear cell and homogenate collection

Mice were anesthetized with isoflurane and perfused with 1x PBS. The brain was removed from the skull, and the spinal cord was flushed through the spinal canal with PBS. Combined brain and spinal cord tissues were homogenized with an 18-gauge needle in a protease inhibitor solution created using protease inhibitor cocktail tablets (Roche) per the manufacturer’s protocol, centrifuged at 800 g for 5 minutes, and the supernatant was collected and stored at −80°C. Tissue pellet was resuspended in a solution of HBSS with 1 mg/ml collagenase A (Roche) and 1 mg/ml DNase I (Sigma-Aldrich) and incubated at 37°C for 20 minutes. Mononuclear cells were separated from myelin via centrifugation in a 27% Percoll solution (GE Healthcare).

### Fluorescent immunohistochemistry

Spinal cords were harvested from mice perfused with 1x PBS and 4% paraformaldehyde (PFA). Tissue were post-fixed with 4% PFA for 24 hours, washed with 1x PBS, decalcified with 0.5 M EDTA for 4–6 days, and cryopreserved with 30% sucrose solution at 4°C. Tissue was embedded in OCT for cryosectioning and stored at −80°C. Sections were blocked with 5% normal goat serum (Sigma-Aldrich), washed with 0.1% Triton-X 100 (Fisher Scientific) in 1x PBS (PBS-T), and stained with rat anti-mouse CD45 (clone IBL-5/25; EMD Millipore) for 48 hours at 4°C. Following washing with PBS-T, sections were stained with Alexa Fluor 647 goat anti-rat IgG (Invitrogen) for 2 hours. Sections were then stained with FluoroMyelin Red Fluorescent Myelin Stain (Invitrogen) per the manufacturer’s protocol. ProLong Gold antifade reagent with DAPI (Invitrogen) was applied immediately prior to applying the coverslip. Images were acquired using an Olympus IX83 inverted fluorescent microscope with cellSens Dimension software (Olympus).

### Electrophysiology

Electrophysiological recordings were performed using a clinical electrodiagnostic system (Cadwell). During recordings mice were anesthetized using ketamine/xylazine anesthesia. A petroleum-based eye lubricant (Dechra) was applied to prevent corneal irritation and dryness. A thermostatically-controlled, far infrared heating pad was used to maintain body temperature (Kent Scientific). Hair from the right hindlimb was removed with clippers (Remington, model VPG 6530) to allow for adequate electrode-skin contact. During anesthetized recordings, mice were placed in the prone position, and bilateral hindlimbs were extended and affixed to the heating pad with Transpore medical tape (3M). Compound muscle action potential (CMAP) and single motor unit potential (SMUP) amplitudes were recorded from the right gastrocnemius muscle similar to our prior studies in aged mice(43, 44). Briefly, a pair of recording ring electrodes (Alpine Biomed) were placed at the proximal gastrocnemius (G1) and at the mid-tarsal region of the hind paw (G2). The skin underneath the ring electrodes was coated with electrode gel (Parker Laboratories) to reduce skin impedance. A disposable surface disc electrode was placed on the surface of the skin of the tail (Natus Neurology) as the common reference electrode (G0). Two 28-gauge monopolar electrodes (Natus Neurology) were inserted subcutaneously at the proximal right thigh and used as the cathode and anode to stimulate the sciatic nerve. A constant current stimulator was used to deliver pulses (0-10 mA current, 0.1 ms duration). Supramaximal stimulation was delivered to record the maximum CMAP amplitude. Then the stimulus was reduced, and a gradually increasing stimulus was used to elicit a total of 10 all-or-none CMAP increments. The 10 incremental responses were averaged to determine the average SMUP amplitude which was used to calculate motor unit number estimation (MUNE = maximum CMAP / Average SMUP). To determine CMAP response following spinal cord conduction, the stimulating electrodes were placed subcutaneously at the base of the skull on each side of the spinal column. The spinal cord was stimulated using a constant current stimulator (0-40 mA, 0.2 ms) to elicit the maximum cervical motor evoked potential (MEP) amplitude. During all recordings, high and low frequency filter settings were set at 10 kHz and 10 Hz, respectively. Peak-to-peak amplitudes were used for all analyses.

### Flow cytometry

For surface staining, cells were resuspended in 1x PBS with 2% fetal bovine serum (FBS), fixable viability dye (eFluor506, eBioscience), and anti-CD16/32 (clone 2.4G2, hybridoma, ATCC). Cells were then labeled with fluorescently labeled monoclonal antibodies specific for individual markers. For intracellular cytokine staining, cells were stimulated with PMA (50 ng/ml), ionomycin (2 μg/ml), and BFA (5 μg/ml) for 4-6 hours. Cells were then fixed and permeabilized with the Fixation/Permeablization Kit (BD Biosciences) according to the manufacturer’s protocols and incubated with fluorescently labeled monoclonal antibodies. Data were acquired using a FACSMelody flow cytometer (BD Biosciences) and analyzed with FlowJo software (Tree Star).

### Antibodies

The following antibodies were obtained from eBiosciences: CD11b (M1/70)- APC-eFluor 780, CD11c (N418)- PerCP-Cyanine5.5, CD45.1 (A20)- FITC and PE, CD45R/B220 (RA3-6B2)- PE, and Ki-67 (SolA15)- PE-Cyanine7. The following antibodies were obtained from Invitrogen: CD45 (30-F11)- eFluor 450 and GM-CSF (MP1-22E9)- FITC. The following antibodies were obtained from BD Pharmingen: Ly6G (1A8)- PE-Cy7. The following antibodies were obtained from BioLegend: CD4 (RM4-5)- APC. For *in vivo* GM-CSF blocking experiments, 500 μg of anti-mouse GM-CSF (MP1-22E9) or rat IgG2a isotype control (2A3) (Bio X Cell) were administered via i.p. injection.

### Multiplex cytokine assay

Cytokine levels were measured in spinal cord homogenates by Luminex multiplex bead-based analysis (MILLIPLEX MAP Mouse Cytokine/Chemokine Magnetic Bead Panel; Millipore) using the Bio-Plex 200 system (Bio-Rad Laboratories) according to the manufacturer’s protocols.

### Generation of bone marrow chimeras

Femur, tibia, and humorous bones were harvested from donor mice. Marrow was flushed from the bone, passed through a 70 μm mesh filter to generate a single-cell suspension, and red blood cells were lysed by a brief incubation in ACK lysis buffer (Quality Biological). Hosts were lethally irradiated with 2 doses of 6.5 Gy and reconstituted by tail vein injection of 2-4 x 10^6^ donor bone marrow cells. Recipient mice were allowed to reconstitute for 6 weeks prior to use.

### Schematic design

All figure schematics were created using BioRender software.

### Single-cell RNAseq analysis

Sequence mapping and preprocessing: Mononuclear cells were isolated from 4 spinal cords per group and processed following the 10X Genomics Chromium Single Cell RNA v3 protocol. Libraries were sequenced on an Illumina Novaseq instrument and counted with CellRanger v3.1.0 using mm10 reference genome GENCODE vM23/Ensembl 98). Data processing and visualizations of the scRNAseq data were performed using the Seurat package (v.4.0.4) in R (4.1.0). For the initial quality control filtering, we removed individual cells that detected less than 200 genes or more than 25,000 reads, and genes that were detected in less than three cells. We filtered outlier cells outside the range of 5x median absolute deviation of that cell due to sequencing depth using scater. We also only retained cells with less than 10% mitochondrial reads and less than 50% ribosomal reads. Data were scaled to 10,000 transcripts per cell, and transformed to log space using Seurat’s LogNormalize method. The top 2,000 highly variable genes in each sample were computed based on dispersion and mean.

#### Data integration and cell clustering

To correct batch effects, anchor genes were identified from all samples using the *FindIntegrationAnchors()* function in Seurat on the first 20 dimensions, and data were integrated via the Seurat *IntegrateData()* function. Principal component analysis (PCA) was performed on the top highly variable genes. The top 20 PCs were used to build a k-nearest-neighbors cell-cell graph with k = 30 neighbors. Cell clusters were identified using the Louvain graph-clustering algorithm with a resolution set to 0.4. We assigned the cell-type labels for each cell cluster using small sets of known marker genes and cluster-specific genes. Finally, the dataset was projected onto two-dimensional space using uniform manifold approximation and projection (UMAP) dimensionality reduction with default parameters.

#### DEG analysis and enrichment test

We identified cluster-specific genes for the clusters using the *FindAllMarkers* function, comparing the gene expression levels in a given cluster with the rest of the cells. The significance of difference was determined using a Wilcoxon Rank Sum test with Bonferroni correction. Differentially expressed genes (DEG) are determined by setting a threshold of the adjusted p-value of < 0.05 and absolute log2-fold change > 0.25. We computationally selected individual cell types for between-group differential gene expression analysis. Note that the batch corrected integration data was only used for cell clustering and dimensional reduction, and the differential gene expression analysis was performed using the normalized RNA assay slot in Seurat. Pathway enrichment analysis was performed using Enrichr using libraries including GO_2018 and KEGG_2019_MOUSE. We also used clusterProfiler’s universal enrichment analyzer to analyze our manually selected GO pathways and overlapping DEGs with public datasets. Volcano plots were generated for DEG visualization using EnhancedVolcano. Heatmaps were generated using ComplexHeatmap on z-score transformed normalized gene expression values.

#### Data availability

Single-cell RNA-seq data will be available in the Gene Expression Omnibus (GEO) database upon acceptance of the manuscript.

### Bulk RNA-seq analysis

Individual FASTQ files were trimmed for adapter sequences and filtered using fastp v0.20.0. Mouse reference genome GRCm38.p6 and gene annotation described by Gene Transfer Format (GTF) were downloaded from Ensembl release 99 (January 2020). Reads alignment was performed against the reference genome using HISAT2 v2.1.0. Gene expression values for genes were quantified using the featureCounts tool of the Subread package v1.5.0-p2 in the unstranded mode. Genes detected in less than ten total counts in at least three samples were removed from downstream analysis. The fold-change was calculated using CPM-normalized values using EdgeR.

### Statistical analysis of clinical, proteomics and flow cytometric data

Statistical analysis was performed in GraphPad Prism using paired or unpaired 2-tailed Student’s t test, or 1-way or 2-way ANOVA with correction for multiple comparisons, as indicated in the legends. Disease curves were compared by unpaired t tests with a Welch correction at individual days where indicated. Comparison for incidence, survival, and remission curves were performed using a Log-rank test. Outliers were identified by ROUT analysis and removed as necessary. A p value less than 0.05 (*) was considered significant; **p <0.01, ***p <0.001, and ****p <0.0001. Statistical analysis used in RNAseq analysis is provided in the previous subsection.

### Study approval

All animal experiments were performed in accordance with an IACUC-approved protocol at The Ohio State University.

## Author Contributions

J.R.A., A.D.J., A.R.S., A. Munie, and W.D.A. performed experiments and data analysis. A. Ma and C.W. oversaw RNAseq analysis. B.M.S. and J.R.A. wrote the manuscript and coedited it with the help of the other authors. B.M.S. and J.R.A. directed the studies.

## Acknowledgements

We thank R. Yousef and A. McVey Moffatt for technical support. This work was supported by National Institute of Neurological Disorders and Stroke grant R01 NS105385 (to BMS). BMS is the Stanley and Joan D. Ross Professor of Neuromodulation.

---

**Supplementary Figure 1.**
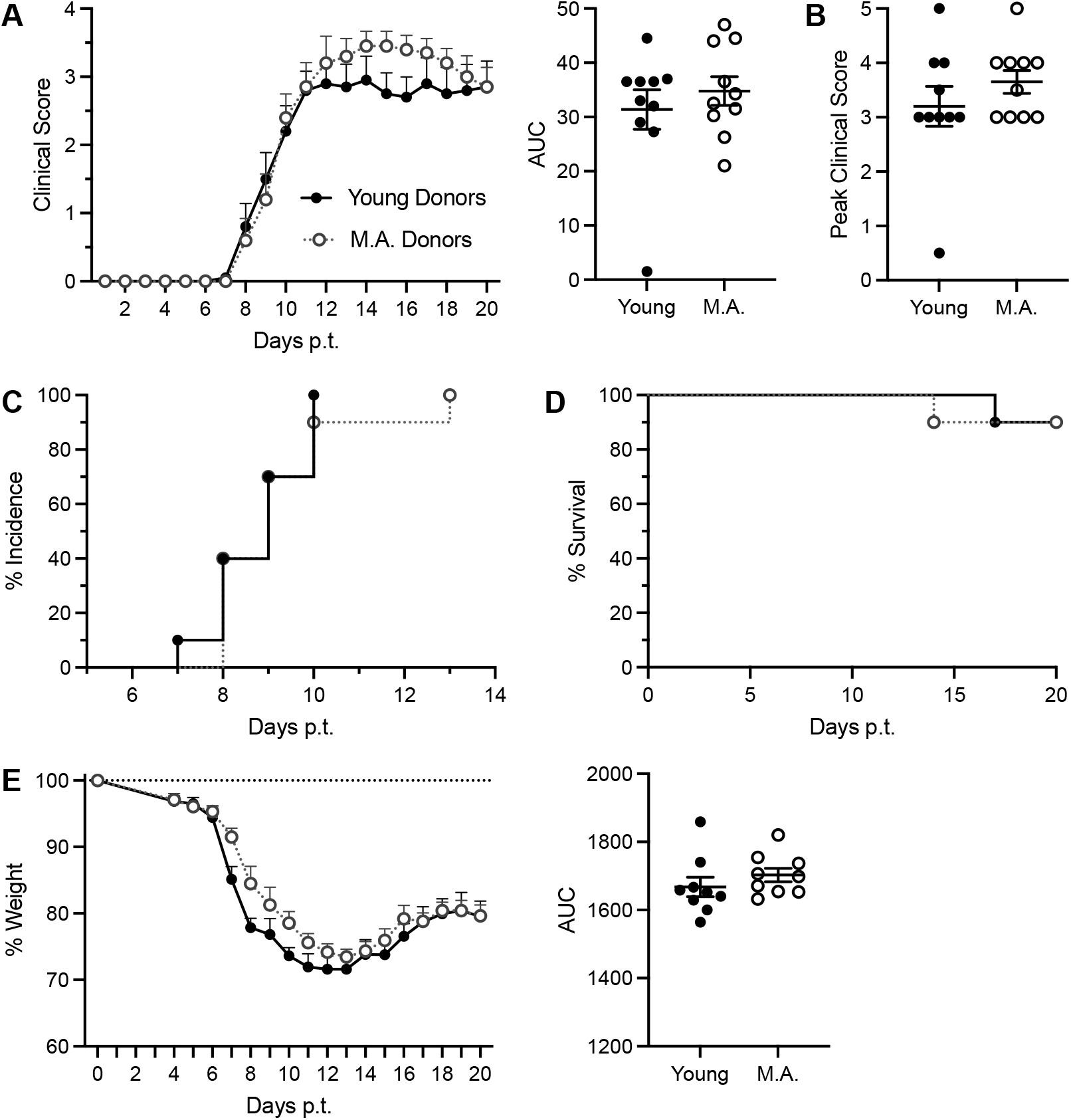
The age of MOG-primed donor CD4+ T cells does not impact the clinical course of adoptively transferred EAE. MOG/CFA primed lymph node cells from young adult (8-12 week old) or middle-age (M.A., 40-44 week old) mice were cultured with MOG_35-55_ peptide and Th17 polarizing factors. The cells were harvested 96 hours later. CD4+ T cells were purified and injected into naïve, syngeneic young adult mice (n=10 mice/ group). (**A**) Mean clinical scores of mice in each group (left) and areas under the curve (AUC) of individual mice (right). (**B**) Peak clinical scores for individual mice. (**C**) Disease incidence in each group over time. (**E**) Percent of surviving mice in each group over time. (**E**) Mean change in weight (shown as percent of baseline weight) of mice in each group throughout the clinical course (left), and AUCs of individual mice (right).

**Supplementary Figure 2.**
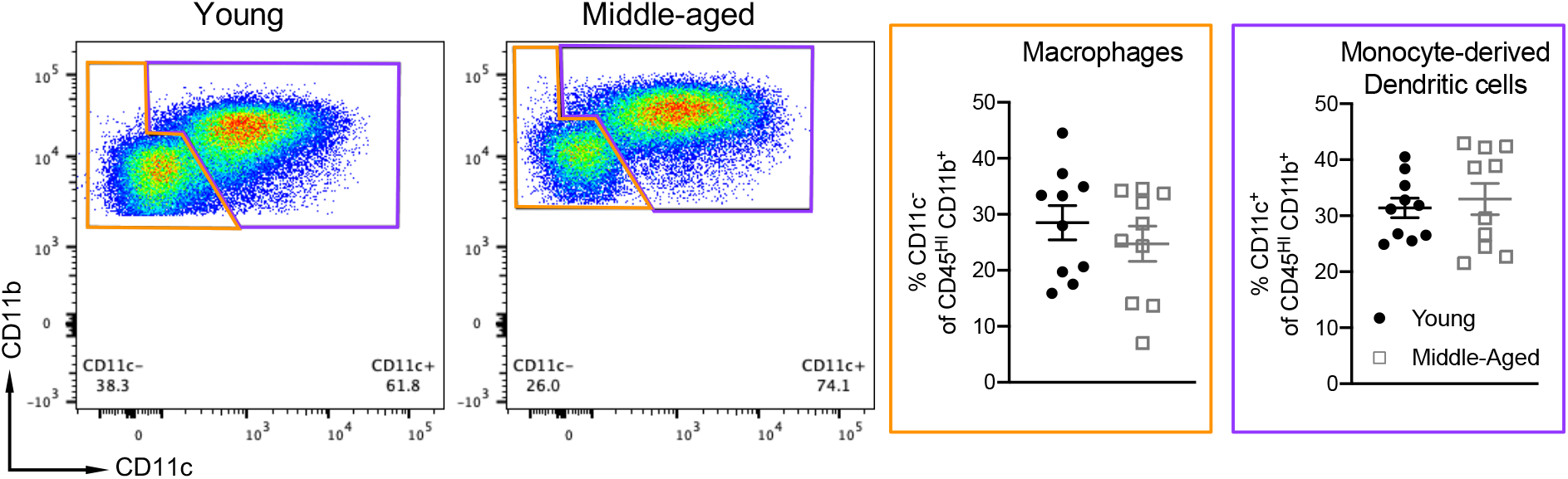
The frequencies of monocyte-derived dendritic cells and macrophages in CNS infiltrates are comparable between young and middle-aged adoptive transfer recipients at peak EAE. CNS inflammatory cells were harvested from the spinal cords of young and middle-aged mice at day 10 adoptive post-transfer. Representative plots (left panels), and frequencies of macrophages (CD45^+^ CD11b^+^ Ly6G^-^ CD11c^-^) and monocyte-derived dendritic cells (CD45^+^ CD11b^+^ Ly6G^-^ CD11c^-^) of individual mice in each group. Data were obtained from n=10 mice/ group pooled from 3 independent experiments. Each symbol represents a data point generated from a single mouse. Statistical significance was determined using the unpaired 2-tailed Student’s *t* test.

**Supplementary Figure 3.**
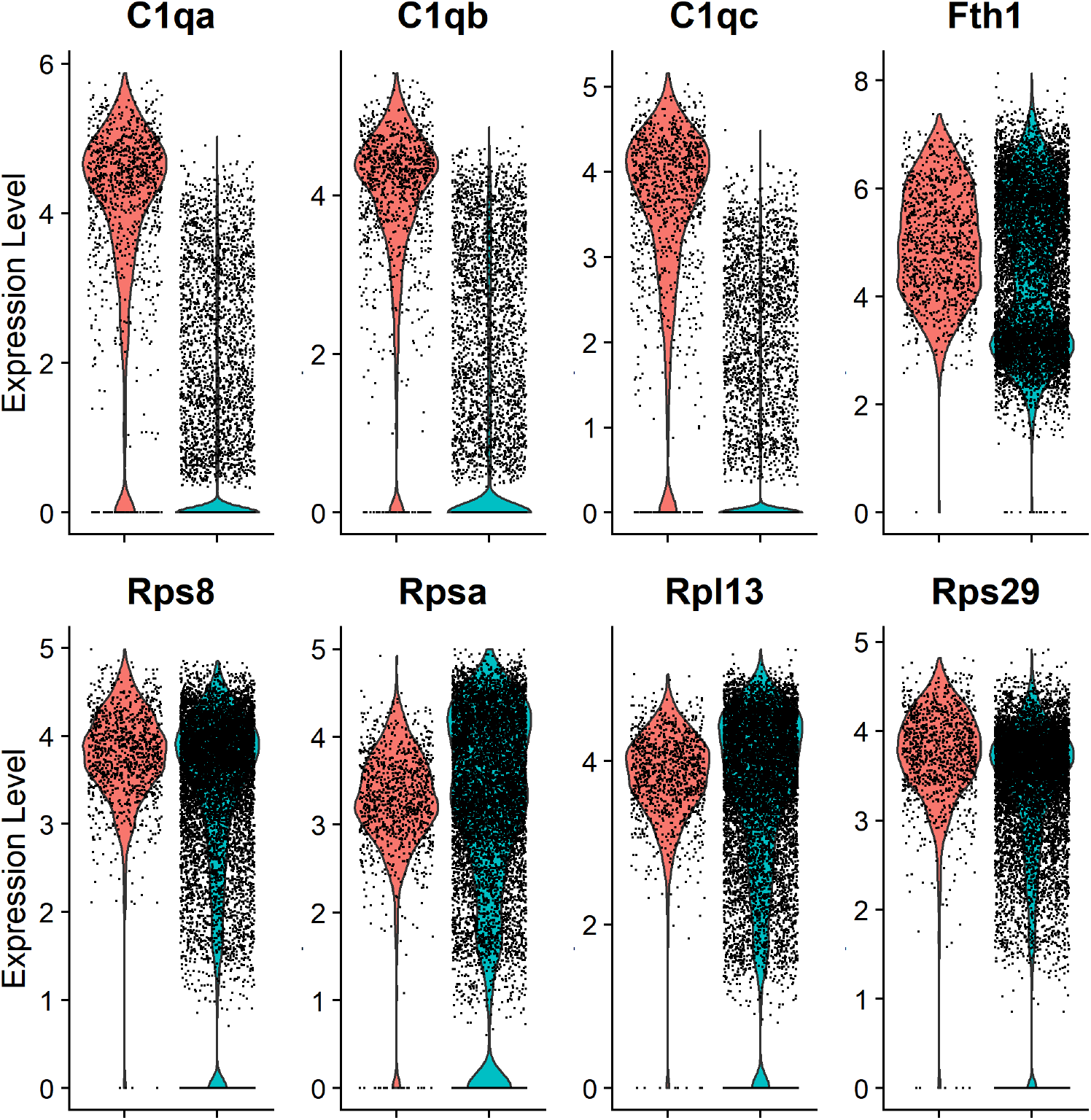
Microglia from middle-aged mice with EAE express high levels of genes identified in microglia located in the rims of chronic active pMS lesions. CD45^+^ mononuclear cells were FACS sorted from middle-aged mice with EAE on day 10 post-cell transfer and subjected to scRNA-seq. Selected differentially expressed genes in microglia (red) compared with all other cells (blue), shown as violin pots. Numbers on the y-axis correspond to the z score for each gene.

## References

1. Lassmann H. Pathogenic Mechanisms Associated With Different Clinical Courses of Multiple Sclerosis. Front Immunol. 2018;9:3116.

2. Frischer JM, Weigand SD, Guo Y, Kale N, Parisi JE, Pirko I, et al. Clinical and pathological insights into the dynamic nature of the white matter multiple sclerosis plaque. Ann Neurol. 2015;78(5):710–21.

3. Tutuncu M, Tang J, Zeid NA, Kale N, Crusan DJ, Atkinson EJ, et al. Onset of progressive phase is an age-dependent clinical milestone in multiple sclerosis. Mult Scler. 2013;19(2):188–98.

4. Harding KE, Liang K, Cossburn MD, Ingram G, Hirst CL, Pickersgill TP, et al. Long-term outcome of paediatric-onset multiple sclerosis: a population-based study. J Neurol Neurosurg Psychiatry. 2013;84(2):141–7.

5. Tremlett H, Zhao Y, Joseph J, Devonshire V, and Neurologists UC. Relapses in multiple sclerosis are age- and time-dependent. J Neurol Neurosurg Psychiatry. 2008;79(12): 1368–74.

6. Rao P, and Segal BM. Experimental autoimmune encephalomyelitis. Methods Mol Med. 2004;102:363–75.

7. Segal BM. CNS chemokines, cytokines, and dendritic cells in autoimmune demyelination. J Neurol Sci. 2005;228(2):210–4.

8. Dutta S, and Sengupta P. Men and mice: Relating their ages. Life Sci. 2016;152:244–8.

9. Lalor SJ, and Segal BM. Th1-mediated experimental autoimmune encephalomyelitis is CXCR3 independent. Eur J Immunol. 2013;43(11):2866–74.

10. Hamann I, Zipp F, and Infante-Duarte C. Therapeutic targeting of chemokine signaling in Multiple Sclerosis. Journal of the Neurological Sciences. 2008;274(1-2):31–8.

11. Carlson T, Kroenke M, Rao P, Lane TE, and Segal B. The Th17-ELR+ CXC chemokine pathway is essential for the development of central nervous system autoimmune disease. J Exp Med. 2008;205(4):811–23.

12. Kroenke MA, Chensue SW, and Segal BM. EAE mediated by a non-IFN-gamma/non-IL-17 pathway. Eur J Immunol. 2010;40(8):2340–8.

13. Duncker PC, Stoolman JS, Huber AK, and Segal BM. GM-CSF Promotes Chronic Disability in Experimental Autoimmune Encephalomyelitis by Altering the Composition of Central Nervous System-Infiltrating Cells, but Is Dispensable for Disease Induction. J Immunol. 2018;200(3):966–73.

14. Hammond TR, Dufort C, Dissing-Olesen L, Giera S, Young A, Wysoker A, et al. Single-Cell RNA Sequencing of Microglia throughout the Mouse Lifespan and in the Injured Brain Reveals Complex Cell-State Changes. Immunity. 2019;50(1):253–71 e6.

15. Mrdjen D, Pavlovic A, Hartmann FJ, Schreiner B, Utz SG, Leung BP, et al. High-Dimensional Single-Cell Mapping of Central Nervous System Immune Cells Reveals Distinct Myeloid Subsets in Health, Aging, and Disease. Immunity. 2018;48(2):380–95 e6.

16. Wynne AM, Henry CJ, and Godbout JP. Immune and behavioral consequences of microglial reactivity in the aged brain. Integr Comp Biol. 2009;49(3):254–66.

17. Perry VH, and Holmes C. Microglial priming in neurodegenerative disease. Nat Rev Neurol. 2014;10(4):217–24.

18. von Bernhardi R, Eugenin-von Bernhardi L, and Eugenin J. Microglial cell dysregulation in brain aging and neurodegeneration. Front Aging Neurosci. 2015;7:124.

19. Batterman KV, Cabrera PE, Moore TL, and Rosene DL. T Cells Actively Infiltrate the White Matter of the Aging Monkey Brain in Relation to Increased Microglial Reactivity and Cognitive Decline. Front Immunol. 2021;12:607691.

20. Gyoneva S, Hosur R, Gosselin D, Zhang B, Ouyang Z, Cotleur AC, et al. Cx3cr1-deficient microglia exhibit a premature aging transcriptome. Life Sci Alliance. 2019;2(6).

21. Manich G, Recasens M, Valente T, Almolda B, Gonzalez B, and Castellano B. Role of the CD200-CD200R Axis During Homeostasis and Neuroinflammation. Neuroscience. 2019;405:118–36.

22. Wynne AM, Henry CJ, Huang Y, Cleland A, and Godbout JP. Protracted downregulation of CX3CR1 on microglia of aged mice after lipopolysaccharide challenge. Brain Behav Immun. 2010;24(7):1190–201.

23. Manabe T, Racz I, Schwartz S, Oberle L, Santarelli F, Emmrich JV, et al. Systemic inflammation induced the delayed reduction of excitatory synapses in the CA3 during ageing. J Neurochem. 2021;159(3):525–42.

24. Huber AK, Giles DA, Segal BM, and Irani DN. An emerging role for eotaxins in neurodegenerative disease. Clin Immunol. 2018;189:29–33.

25. Huber AK, Wang L, Han P, Zhang X, Ekholm S, Srinivasan A, et al. Dysregulation of the IL-23/IL-17 axis and myeloid factors in secondary progressive MS. Neurology. 2014;83(17):1500–7.

26. Haghikia A, Jorg S, Duscha A, Berg J, Manzel A, Waschbisch A, et al. Dietary Fatty Acids Directly Impact Central Nervous System Autoimmunity via the Small Intestine. Immunity. 2015;43(4):817–29.

27. Rothhammer V, Borucki DM, Tjon EC, Takenaka MC, Chao CC, Ardura-Fabregat A, et al. Microglial control of astrocytes in response to microbial metabolites. Nature. 2018;557(7707):724–8.

28. Duscha A, Gisevius B, Hirschberg S, Yissachar N, Stangl GI, Eilers E, et al. Propionic Acid Shapes the Multiple Sclerosis Disease Course by an Immunomodulatory Mechanism. Cell. 2020;180(6):1067–80 e16.

29. Ruiz-Ruiz S, Sanchez-Carrillo S, Ciordia S, Mena MC, Mendez-Garcia C, Rojo D, et al. Functional microbiome deficits associated with ageing: Chronological age threshold. Aging Cell. 2020;19(1):e13063.

30. Mockus TE, Munie A, Atkinson JR, and Segal BM. Encephalitogenic and Regulatory CD8 T Cells in Multiple Sclerosis and Its Animal Models. J Immunol. 2021;206(1):3–10.

31. Booss J, Esiri MM, Tourtellotte WW, and Mason DY. Immunohistological analysis of T lymphocyte subsets in the central nervous system in chronic progressive multiple sclerosis. J Neurol Sci. 1983;62(1-3):219–32.

32. Machado-Santos J, Saji E, Troscher AR, Paunovic M, Liblau R, Gabriely G, et al. The compartmentalized inflammatory response in the multiple sclerosis brain is composed of tissue-resident CD8+ T lymphocytes and B cells. Brain. 2018;141(7):2066–82.

33. Stoolman JS, Duncker PC, Huber AK, Giles DA, Washnock-Schmid JM, Soulika AM, et al. An IFNgamma/CXCL2 regulatory pathway determines lesion localization during EAE. J Neuroinflammation. 2018;15(1):208.

34. Stoolman JS, Duncker PC, Huber AK, and Segal BM. Site-specific chemokine expression regulates central nervous system inflammation and determines clinical phenotype in autoimmune encephalomyelitis. J Immunol. 2014;193(2):564–70.

35. Balashov KE, Rottman JB, Weiner HL, and Hancock WW. CCR5(+) and CXCR3(+) T cells are increased in multiple sclerosis and their ligands MIP-1alpha and IP-10 are expressed in demyelinating brain lesions. Proc Natl Acad Sci U S A. 1999;96(12):6873–8.

36. Simpson JE, Newcombe J, Cuzner ML, and Woodroofe MN. Expression of the interferon-gamma-inducible chemokines IP-10 and Mig and their receptor, CXCR3, in multiple sclerosis lesions. Neuropathol Appl Neurobiol. 2000;26(2):133–42.

37. Fox RJ, Kivisakk P, Fisher E, Tucky B, Lee JC, Rudick RA, et al. Multiple sclerosis: chemokine receptor expression on circulating lymphocytes in correlation with radiographic measures of tissue injury. Mult Scler. 2008;14(8):1036–43.

38. Marzan DE, Brugger-Verdon V, West BL, Liddelow S, Samanta J, and Salzer JL. Activated microglia drive demyelination via CSF1R signaling. Glia. 2021;69(6):1583–604.

39. Clark KC, Josephson A, Benusa SD, Hartley RK, Baer M, Thummala S, et al. Compromised axon initial segment integrity in EAE is preceded by microglial reactivity and contact. Glia. 2016;64(7):1190–209.

40. Liddelow SA, Guttenplan KA, Clarke LE, Bennett FC, Bohlen CJ, Schirmer L, et al. Neurotoxic reactive astrocytes are induced by activated microglia. Nature. 2017;541(7638):481–7.

41. Calderwood SK, Borges TJ, Eguchi T, Lang BJ, Murshid A, Okusha Y, et al. Extracellular Hsp90 and protection of neuronal cells through Nrf2. Biochem Soc Trans. 2021;49(5):2299–306.

42. Yu WW, Cao SN, Zang CX, Wang L, Yang HY, Bao XQ, et al. Heat shock protein 70 suppresses neuroinflammation induced by alpha-synuclein in astrocytes. Mol Cell Neurosci. 2018;86:58–64.

43. Chugh D, Iyer CC, Wang X, Bobbili P, Rich MM, and Arnold WD. Neuromuscular junction transmission failure is a late phenotype in aging mice. Neurobiol Aging. 2020;86:182–90.

44. Sheth KA, Iyer CC, Wier CG, Crum AE, Bratasz A, Kolb SJ, et al. Muscle strength and size are associated with motor unit connectivity in aged mice. Neurobiol Aging. 2018;67:128–36.

